# Functional analysis of *PRPS1* disease alleles reveals altered polymerization and cytoskeletal organization

**DOI:** 10.64898/2026.10.26.761893

**Authors:** Elena Monfort, Risa M. Broyer, Juan-José Ripoll, James E. Wilhelm

## Abstract

Phosphoribosyl pyrophosphate synthetase 1 catalyzes a central step in nucleotide biosynthesis, and pathogenic variants cause a spectrum of neurological disorders ranging from loss-of-function syndromes to enzyme superactivity. Despite their opposing biochemical effects, these variants often produce overlapping phenotypes, suggesting that shared cellular processes may contribute to disease pathogenesis. Here, using *Saccharomyces cerevisiae* and patient-derived human fibroblasts, we examined how disease-associated variants affect phosphoribosyl pyrophosphate synthetase 1 polymerization and whether changes in assembly are associated with metabolic state and cytoskeletal organization. Human phosphoribosyl pyrophosphate synthetase 1 functionally substituted for essential phosphoribosyl pyrophosphate synthetase paralogs in *Saccharomyces cerevisiae*, and disease-associated variants differentially altered filament abundance, length, and architecture in ways not fully explained by changes in protein levels. Polymer assembly was sensitive to cellular metabolic state: deletion of the yeast adenosine kinase gene, a condition previously shown to elevate intracellular adenosine and S-adenosylmethionine, strongly suppressed filament formation, including in hyperpolymerizing superactivity variants. Both gain- and loss-of-function variants were associated with altered actin organization in *Saccharomyces cerevisiae* and patient-derived human fibroblasts, whereas the D183H superactivity variant promoted aberrant cytoplasmic polymerization in human cells. Together, these findings identify phosphoribosyl pyrophosphate synthetase 1 polymerization as a metabolically responsive feature associated with cytoskeletal organization and provide a cellular framework for understanding how functionally distinct disease alleles can converge on overlapping phenotypes.

## Introduction

Mutations in genes encoding metabolic enzymes can give rise to diverse clinical phenotypes, yet how distinct biochemical perturbations converge on shared cellular outcomes remains poorly understood. Phosphoribosyl pyrophosphate synthetase 1 (PRPS1), a central enzyme in nucleotide biosynthesis, provides a notable example. Pathogenic variants in this X-linked gene span a spectrum from loss-of-function syndromes to enzyme superactivity, yet these conditions frequently share neurological manifestations. This apparent disconnect between biochemical activity and phenotypic outcome suggests that additional cellular processes may contribute to the convergence of PRPS1-associated disease phenotypes.

PRPS1 catalyzes the synthesis of phosphoribosyl pyrophosphate (PRPP), an essential precursor for purine, pyrimidine, NAD, and amino acid biosynthesis. Pathogenic PRPS1 alleles encompass functionally distinct classes. Gain-of-function (GOF) mutations that increase enzyme activity elevate purine production and can lead to hyperuricemia and gout (Becker and Seegmiller 1975; Becker et al. 1988). Loss-of-function (LOF) mutations cause disorders including nonsyndromic sensorineural deafness (DFN2), X-linked Charcot–Marie–Tooth neuropathy type 5 (CMTX5), and Arts syndrome, which variably include hearing loss, ataxia, peripheral neuropathy, and, in severe cases, intellectual disability (Rosenberg and Chutorian 1967; Arts et al. 1993; de Brouwer et al. 2007). A subset of GOF mutations, commonly referred to as superactivity alleles, disrupt feedback inhibition and produce hyperuricemia together with neurological manifestations that overlap those associated with LOF variants (Akaoka et al. 1981; Becker et al. 1980; 1982; 1986; 1988; Becker et al. 1995; 1996; Ahmed et al. 1999). Thus, variation in catalytic activity alone does not readily account for the convergence of neurological phenotypes across functionally distinct PRPS1 alleles, raising the possibility that additional aspects of PRPS1 regulation contribute to disease.

Therapeutic options for PRPS1-related disorders remain limited. Xanthine oxidase inhibitors can control hyperuricemia associated with superactivity, but they do not address the neurological manifestations. Metabolic supplementation has therefore been explored as an alternative strategy. S-adenosylmethionine (SAM) has been proposed to bypass PRPP-dependent purine synthesis, and combined SAM and nicotinamide riboside treatment improved survival and metabolic function in PRPS1-deficient human T cells (Lenherr et al. 2021). These observations underscore the potential importance of metabolic context in PRPS1 dysfunction and motivate a broader understanding of how PRPS1 is regulated within cells.

Previous systematic screens revealed that multiple metabolic enzymes assemble into higher-order filamentous structures, suggesting an additional level of spatial organization in cellular metabolism. Members of the phosphoribosyl pyrophosphate synthetase family, including PRPS1, form reversible filaments in yeast and human cells (Noree et al. 2019), indicating that higher-order assembly is an evolutionarily conserved feature of this enzyme family. However, the functional significance of PRPS1 filament formation and its relationship to disease-associated mutations remain poorly understood. In particular, it remains unclear whether changes in polymerization accompany altered PRPS1 functional states and how polymerization responds to the cellular metabolic environment.

*Saccharomyces cerevisiae* provides a tractable system in which to address these questions because PRPP synthesis and many of its regulatory connections are conserved across eukaryotes. Heterologous expression in yeast has been widely used to investigate human disease genes and pathogenic variants, while comparison with mammalian cells provides a means to distinguish conserved cellular responses from system-specific features (Botstein and Fink 2011). This cross-species approach therefore provides an opportunity to examine the cellular consequences of PRPS1 disease variants in genetically and experimentally complementary systems.

The *S. cerevisiae* genome contains five *PRS* genes whose products, Prs1p–Prs5p, form multimeric complexes required for PRPP synthesis. *PRS1* and *PRS3* have particularly important roles, as their deletion strongly reduces PRPP levels, nucleotide pools, and cell-integrity signaling, whereas loss of *PRS2*, *PRS4*, or *PRS5* produces milder phenotypes (Hove-Jensen et al. 2017). Beyond PRPP synthesis, yeast PRS proteins contribute to cell-wall integrity, ion homeostasis, cell-cycle regulation, and actin cytoskeletal organization and participate in nutrient-responsive growth networks (Binley et al. 1999; Jorgensen et al. 2004; Wang et al. 2004; Ugbogu et al. 2016). Despite this functional conservation, whether human PRPS1 can substitute for specific yeast PRS proteins and how human disease-associated variants influence polymerization and associated cellular phenotypes in yeast have not been systematically examined.

Here, we used a combined yeast–mammalian approach to determine how disease-associated PRPS1 variants affect enzyme function, polymerization, and cellular organization. We first tested whether human PRPS1 can functionally substitute for endogenous yeast PRS proteins and then examined a panel of clinically relevant variants for effects on complementation, filament formation, protein abundance, and cellular phenotypes. We further asked whether PRPS1 assembly is sensitive to metabolic context and whether related polymerization and cytoskeletal phenotypes are observed in patient-derived fibroblasts. Together, these experiments provide a cross-species framework for examining how functionally distinct PRPS1 disease alleles alter higher-order enzyme assembly and converge on shared cellular phenotypes.

## Materials and Methods

### Yeast strains and growth media

The *Saccharomyces cerevisiae* strain YER105C from the Yeast GFP Clone Collection (MATa *his3*Δ*1 leu2*Δ*0 met15*Δ*0 ura3*Δ*0 Nup157::GFP*) was used as the background strain for all yeast experiments. A list of human PRPS1 variants examined in this study is provided in Table S1.

Centromeric ARS/CEN plasmids are maintained at low copy number and exhibit high mitotic stability in *S. cerevisiae* (Murray and Szostak 1983; Hieter et al. 1985; Sikorski and Hieter 1989). Because auxotrophic selection markers can influence relative growth behavior and cellular physiology (Karim et al. 2013), strains were grown in YPD medium (2% peptone, 1% yeast extract, 2% dextrose) for liquid-culture growth and phenotypic analyses unless otherwise indicated. Cultures were inoculated from freshly streaked selective plates, grown overnight at 30 °C, diluted to OD₆₀₀ = 0.2, and harvested at mid-log phase unless otherwise noted.

### Deletion mutant construction

Single-gene knockouts of *PRS1* and *PRS3* were generated in the YER105C background using the lithium acetate/PEG transformation method (Ito et al. 1983). PCR-generated deletion cassettes carrying either NAT or *kanMX6* resistance markers were amplified using primers containing 40-bp homology regions corresponding to the 5′ and 3′ regions of each target locus. Transformants were selected on antibiotic-containing YPD plates.

For the complementation assay, *PRS5* was disrupted in *prs3*Δ strains expressing wild-type or disease-associated human PRPS1 by transformation with a PCR-generated deletion cassette carrying the NAT nourseothricin-resistance marker. *ADO1* was similarly disrupted in the Nup157– GFP *prs3*Δ background by transformation with a PCR-generated NAT resistance cassette to generate the *prs3*Δ *ado1*Δ strain used for the metabolic-context experiments. All gene disruptions were confirmed by diagnostic PCR.

### *PRPS1* cloning

The full-length human *PRPS1* coding sequence was amplified from clone HsCD00041572 (DNASU Plasmid Repository). Primers introducing XbaI and XhoI restriction sites were used for directional cloning into the low-copy CEN6/ARS4 plasmid pRH1939 (gift from R. Hampton, UCSD), a derivative of p416-GPD (Mumberg et al. 1995) in which expression is driven by the constitutive *TDH3* promoter. Plasmids were transformed into yeast and transformants were selected on SC–Ura medium.

Disease-associated *PRPS1* variants were generated in pRH1939-*PRPS1* by PCR-based site-directed mutagenesis using allele-specific primers. All constructs were verified by Sanger sequencing.

### Serial spot assay

A single SC–Ura colony was inoculated into 5 mL YPD and grown overnight at 30 °C. Cultures were adjusted to OD_600_ = 0.5, serially diluted 10-fold, and 3 µL of each dilution were spotted onto SC–Ura plates to maintain plasmid selection. Plates were incubated at 30 °C for 48 h before imaging.

### Immunostaining and fluorescence confocal microscopy

For yeast, indirect immunofluorescence was performed as described previously (Noree et al. 2010). Cells were incubated overnight at 4 °C with rabbit anti-PRPS1 antibody (1:1000; Noree et al. 2019) and/or mouse anti-actin antibody (JLA20; 1:100; DSHB), followed by Alexa Fluor 568– conjugated anti-rabbit and/or anti-mouse secondary antibodies (1:200; Invitrogen) for 2 h at room temperature.

Images were acquired as multi-plane Z-stacks using a Zeiss Axiovert 200M microscope equipped with a CSU-X1 spinning-disk confocal unit (Yokogawa), an iChrome MLE laser source (Toptica Photonics), and a 100×/1.4 NA oil-immersion objective.

Primary human fibroblasts were cultured on glass coverslips in MEM lacking glutamine (Gibco) supplemented with 10% fetal bovine serum, L-glutamine, and penicillin–streptomycin. Cell lines WT, D183H, LOF1, and LOF2 were used in this study. The D183H fibroblast line, carrying the *PRPS1* superactivity variant, was established from the patient originally described by Nyhan et al. (1969) during clinical evaluation for hyperuricemia and behavioral abnormalities and was subsequently maintained as an archived fibroblast line. Cells were fixed in 4% paraformaldehyde in PBS for 10 min at room temperature, permeabilized in PBS containing 0.5% Triton X-100 and 1% goat serum, and incubated overnight at 4 °C with primary antibodies diluted in PBS containing 1% goat serum. After washing, coverslips were incubated with Alexa Fluor 568–conjugated anti-rabbit secondary antibody for 2 h at room temperature. Filamentous actin was visualized using rhodamine–phalloidin (1:200; Thermo Fisher Scientific).

Confocal images were acquired using a Leica TCS SP5 microscope with a 63× objective, collecting full Z-stacks at 0.8 µm intervals. Image-acquisition settings were kept identical across conditions, and fields of view were selected in an unbiased manner.

### Image quantification and morphometric analysis

Image processing and quantitative analyses were performed in FIJI (ImageJ). PRPS1 filament length was measured manually using the segmented-line tool on maximum-intensity projections of confocal Z-stacks. Only discrete, continuous PRPS1-positive structures clearly distinguishable from diffuse cytoplasmic signal and ≥0.5 µm in length were included in the analysis. These criteria were applied uniformly across all samples. For filament-length analyses, 94 filaments per strain were analyzed across three independent biological replicates.

For analysis of yeast actin organization, more than 100 cells per strain were scored in each biological replicate for the presence of dense F-actin aggregates. Additional actin abnormalities, including elongated cable-like structures, non-polarized cortical patches, transverse actin bundles, and abnormal accumulation near the mother–bud neck, were evaluated qualitatively because of their morphological heterogeneity. Imaging analyses were performed in at least three independent biological replicates.

For cell-size and morphology analyses, only unbudded cells were included to avoid confounding effects of bud emergence. Cell segmentation and morphometric measurements were performed using identical thresholding parameters across all strains. Cell circularity was calculated as 4π × area/perimeter², with values approaching 1 indicating a more circular cell shape. Cell-size distributions were additionally visualized by kernel density estimation.

Actin stress-fiber orientation in human fibroblasts was quantified using Local Gradient Analysis in FIJI. Quantitative analyses were performed using predefined criteria applied uniformly across samples.

### Statistical analysis

Quantitative data are presented as mean ± SD where indicated. Filament-length distributions were compared separately for gain-of-function and loss-of-function PRPS1 variants using two-sided Mann–Whitney U tests, with each variant compared with wild-type PRPS1.

For analysis of yeast cell circularity, statistical significance relative to wild-type cells was assessed using a Kruskal–Wallis test followed by Dunn’s multiple-comparison test. For analyses examining the relationship between PRPS1 protein abundance and filament frequency or filament length, the coefficient of determination (*R*²) is reported. Biological replicate numbers and sample sizes are provided in the corresponding figure legends.

The frequency of cells containing F-actin aggregates was compared between *prs3*Δ and each PRPS1-expressing strain using two-sided unpaired Student’s *t*-tests.

### Protein extraction and immunoblotting

For yeast, whole-cell extracts were prepared as described previously (Noree et al. 2019) and analyzed by SDS–PAGE and immunoblotting. PRPS1 abundance was quantified relative to tubulin and normalized to wild-type PRPS1 levels. Representative immunoblots shown in Figure S2 derive from the same experimental series used for the protein-abundance quantifications presented in Figs. 2b and 3b.

For human fibroblasts, cells were grown to confluency, washed with PBS, and lysed in RIPA buffer supplemented with protease inhibitors (1:1000; Sigma). Lysates were clarified by centrifugation and analyzed by SDS–PAGE and immunoblotting. Actin was used as the loading control for fibroblast extracts.

Primary antibodies were rabbit anti-PRPS1 (1:1000), mouse anti-tubulin (12G10; 1:1000; DSHB), and mouse anti-actin (JLA20; 1:100; DSHB). HRP-conjugated donkey anti-rabbit IgG and HRP-conjugated sheep anti-mouse IgG secondary antibodies were used at 1:10,000 (GE Healthcare).

### Ethics statement

The D183H *PRPS1* fibroblast line was established from the patient originally described by Nyhan et al. (1969) during clinical evaluation for hyperuricemia and behavioral abnormalities, before contemporary institutional review board procedures were established. The cells were subsequently maintained as an archived fibroblast line. No new human participants were recruited and no new patient material was collected for this study; therefore, prospective IRB approval was not required for the work reported here.

## Results

### Human PRPS1 complements the yeast *prs1*Δ background and rescues its slow-growth phenotype

To determine whether human PRPS1 is functional in yeast, we expressed wild-type PRPS1 in a *prs1*Δ strain. Heterologous expression of human PRPS1 restored growth to near wild-type levels in serial dilution assays (Fig. 1a), indicating that PRPS1 can functionally compensate for loss of endogenous Prs1p in yeast. This phenotype was reproducible across biological replicates.

**Figure 1.**
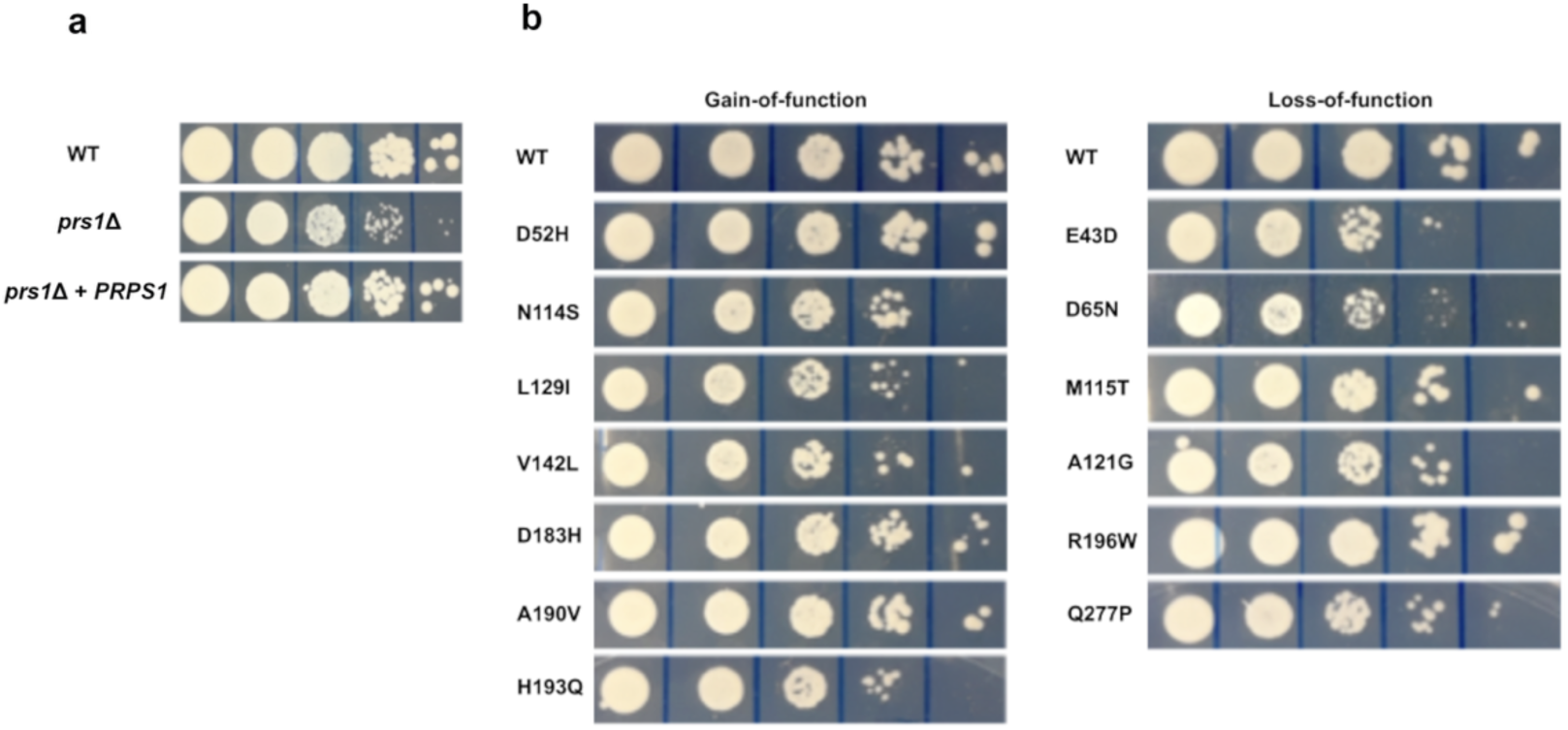
Human PRPS1 supports viability following *PRS5* deletion in a *prs3*Δ background. (a) Serial 10-fold dilution assays showing that heterologous expression of human *PRPS1* rescues the slow-growth phenotype of *prs1*Δ cells. Wild-type (WT), *prs1*Δ, and *prs1*Δ cells expressing human *PRPS1* from a low-copy CEN plasmid were grown on SC–Ura medium for 48 h at 30 °C. (b) Human *PRPS1* supports growth following *PRS5* deletion in a *prs3*Δ background. A *prs3*Δ strain expressing wild-type or disease-associated human *PRPS1* alleles from low-copy CEN plasmids was transformed with a *PRS5* deletion cassette. Growth of the resulting transformants was assessed by serial dilution on SC–Ura medium at 30 °C for 48 h. All strains in panel (b) share the same *prs3*Δ background. For clarity, variant labels are abbreviated in subsequent figures.

### Human PRPS1 supports viability following PRS5 deletion in a *prs3*Δ background

Previous studies reported that simultaneous loss of *PRS3* and *PRS5* is synthetically lethal in yeast (Hernando et al. 1998). To determine whether human PRPS1 could functionally substitute in this context, we expressed plasmid-borne PRPS1 in a *prs3*Δ strain and subsequently disrupted *PRS5*. Wild-type human PRPS1 supported robust growth following PRS5 deletion on SC–Ura medium (Fig. 1b), indicating that human PRPS1 can provide essential PRS function in the absence of both Prs3p and Prs5p.

### Yeast complementation distinguishes functional differences among *PRPS1* disease alleles

Having established functional complementation by wild-type PRPS1, we next evaluated a panel of previously described disease-associated human PRPS1 alleles (Table S1) for their ability to support viability following PRS5 deletion in a *prs3*Δ background. PRPS1 variants were expressed from a centromeric CEN6/ARS4 plasmid (Sikorski and Hieter 1989), which is typically maintained at low copy number in yeast and minimizes the effects associated with high-copy overexpression vectors. Because all alleles were expressed from the same low-copy plasmid under a common promoter, differences in rescue efficiency are consistent with allele-specific functional differences, although variation in steady-state expression among variants cannot be excluded.

Most gain- and loss-of-function alleles supported growth, but with distinct efficiencies (Fig. 1b), revealing a graded spectrum of functional complementation across PRPS1 variants (Table S2). Among gain-of-function variants, D52H and A190V supported growth comparably to wild-type PRPS1, whereas D183H and N114S produced intermediate rescue. V142L, H193Q, and L129I supported weaker growth. Among loss-of-function variants, R196W restored growth to near wild-type levels, followed by M115T. A121G, Q277P, D65N, and E43D conferred weak rescue. Q133P was the only allele unable to support viability following *PRS5* deletion (Fig. 1b).

Together, these results show that yeast complementation can distinguish functional differences among PRPS1 disease alleles at the cellular level.

### Mutational analysis of PRPS1 polymerization in yeast

Having established that human PRPS1 variants differentially support viability in a *prs3*Δ *prs5*Δ background, we next asked whether these functional differences were associated with altered PRPS1 polymerization. PRPS1 variants were expressed in a *prs3*Δ background under identical plasmid and promoter conditions, separately from the PRS5 deletion assay used to assess viability. Under these conditions, wild-type PRPS1 formed distinct cytoplasmic filaments, consistent with previous reports (Noree et al. 2019). Cells were scored as filament-positive if they contained at least one discrete, continuous PRPS1-positive structure that was distinguishable from diffuse cytoplasmic signal and ≥0.5 μm in length, according to the criteria described in Materials and Methods.

Disease-associated variants differentially affected filament frequency and morphology. The feedback-resistant superactivity variants D52H and D183H produced approximately twofold more filament-containing cells than wild-type PRPS1 (Fig. 2a). An independent analysis performed in a wild-type *PRS* background revealed a similar increase in filament frequency for D52H and D183H (Figure S1), indicating that increased filament formation is maintained across distinct yeast genetic backgrounds. In parallel, cells expressing these variants accumulated PRPS1 protein at levels approximately 2–5-fold higher than wild type (Fig. 2b; Figure S2). Because all variants were expressed from low-copy CEN plasmids under a common promoter, differences in plasmid context were minimized, although allele-specific differences in steady-state protein abundance remained evident.

**Figure 2.**
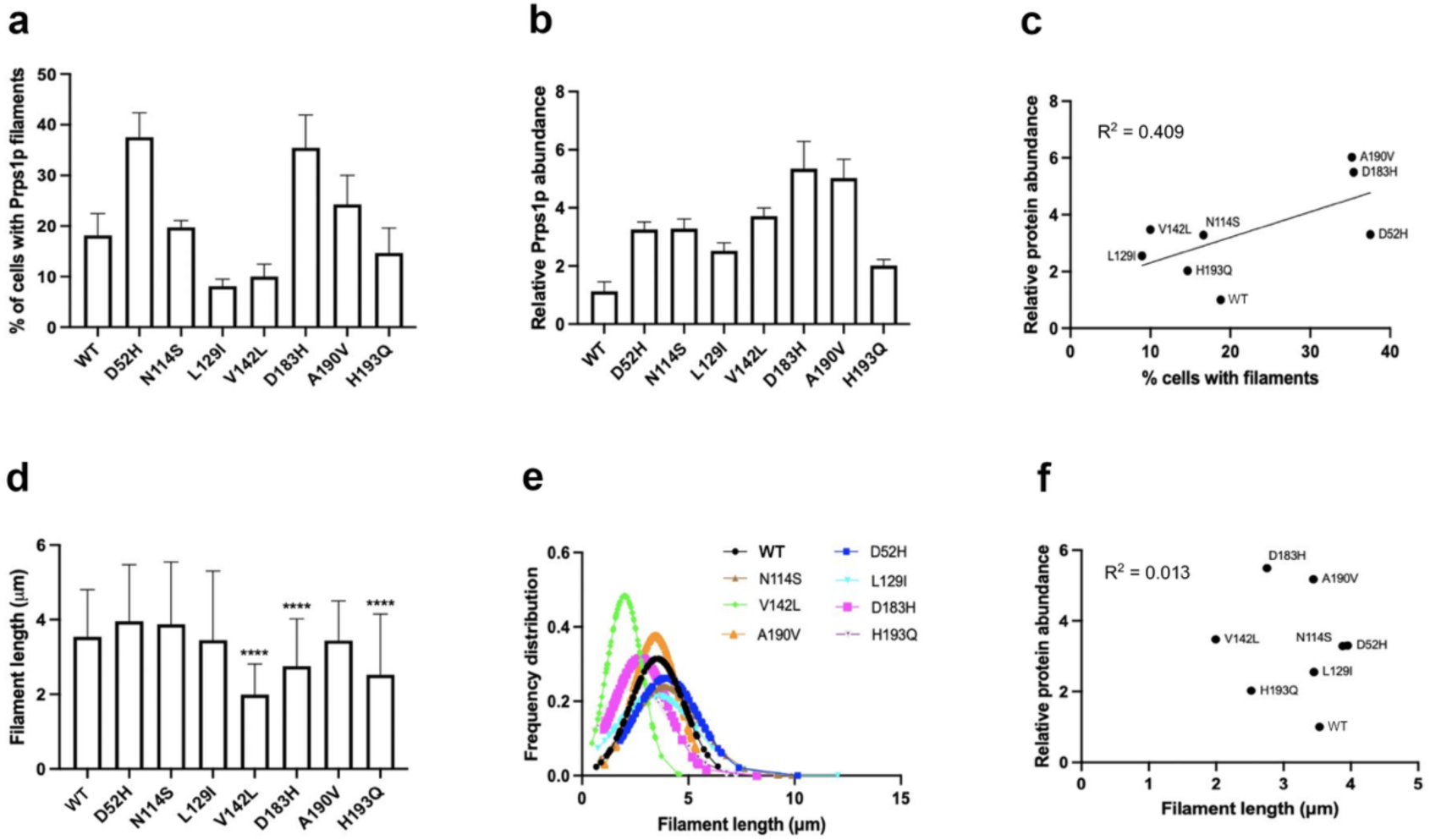
Gain-of-function PRPS1 variants alter filament abundance, protein levels, and filament length in yeast. (a) Percentage of logarithmically growing yeast cells containing PRPS1 filaments. Superactivity-associated variants D52H and D183H exhibited increased filament frequency relative to wild-type PRPS1, whereas L129I and V142L formed filaments at reduced frequencies. (b) Relative PRPS1 protein abundance for the same strains, quantified as PRPS1/tubulin and normalized to wild-type levels. Several GOF variants accumulated at elevated levels relative to wild type. (c) Correlation between filament formation frequency and PRPS1 abundance across GOF variants. A moderate positive correlation was observed (R² = 0.409). (d) Mean filament lengths for wild-type and GOF PRPS1 variants. V142L, D183H, and H193Q formed significantly shorter filaments than wild type, whereas the remaining variants showed no detectable difference. (e) Kernel density distributions of filament lengths for wild-type and GOF variants. Wild-type filaments displayed an approximately normal length distribution, whereas the remaining variants displayed distinct filament-length distributions. (f) Correlation between filament length and PRPS1 protein abundance across GOF variants. No correlation was observed (R² = 0.013). For filament-length measurements, 94 filaments per strain were analyzed across three independent biological replicates. Data are presented as mean ± SD where applicable. Filament-length distributions were compared using two-sided Mann–Whitney U tests comparing each variant with wild-type PRPS1. ****P < 0.0001.

### Additional allosteric-site mutations differentially affect filament organization

Because many gain-of-function (GOF) alleles alter allosteric regulation, we next examined additional variants affecting the two major regulatory sites of PRPS1. Several of these mutations have been reported to impair feedback inhibition or otherwise alter metabolic regulation (Li et al. 2007; Hvorecny et al. 2023).

The GOF variants N114S and H193Q affect residues associated with the allosteric regulatory sites and have been reported to impair feedback inhibition while retaining near-normal kinetic properties (Li et al. 2007; de Brouwer et al. 2010). Both variants formed filaments at frequencies comparable to wild-type PRPS1 (Fig. 2a). In contrast, A190V, which affects allosteric site II without directly altering the catalytic active site, showed increased filament formation and additionally generated sheet-like assemblies in approximately 10% of cells. These sheet-like structures were excluded from filament quantification. This phenotype is consistent with previous structural predictions that A190V perturbs hydrophobic packing at the dimerization interface (Chen et al. 2015).

Conversely, L129I and V142L, both associated with altered allosteric regulation (de Brouwer et al. 2010; Moran et al. 2012), formed approximately half as many filaments as wild-type PRPS1 (Fig. 2a). V142L has previously been described as an intermediate clinical allele exhibiting both gain- and loss-of-function features (Moran et al. 2012). Across the GOF genotype panel, filament frequency and PRPS1 protein abundance showed a moderate positive correlation (R² = 0.409; n = 8 genotypes; Fig. 2c), indicating that protein abundance may contribute to, but does not fully account for, the observed differences in filament formation.

### Filament length is selectively reduced in specific gain-of-function variants

We next quantified filament length for wild-type and GOF PRPS1 variants (Fig. 2d). V142L, D183H, and H193Q produced significantly shorter filaments than wild-type PRPS1, whereas the remaining variants showed no detectable difference. Wild-type filament lengths approximated a normal distribution (Fig. 2e). In contrast to filament frequency, filament length showed no correlation with PRPS1 protein abundance (R² = 0.013; Fig. 2f). Together, these data show that GOF variants differentially affect both filament frequency and filament architecture, and that variation in filament length is not explained by steady-state PRPS1 abundance.

### Loss-of-function *PRPS1* alleles differentially affect filament formation, protein abundance, and filament length

Because GOF variants differentially affected filament organization and protein abundance, we next examined whether LOF alleles produced distinct polymerization phenotypes. LOF PRPS1 alleles are associated with a spectrum of clinically distinct syndromes and exhibit graded differences in functional activity. We therefore examined whether these variants also differed in PRPS1 abundance and polymerization behavior. When expressed under the same plasmid and promoter conditions as wild-type PRPS1, LOF variants exhibited diverse and allele-specific effects on both protein abundance and filament formation (Fig. 3).

**Figure 3.**
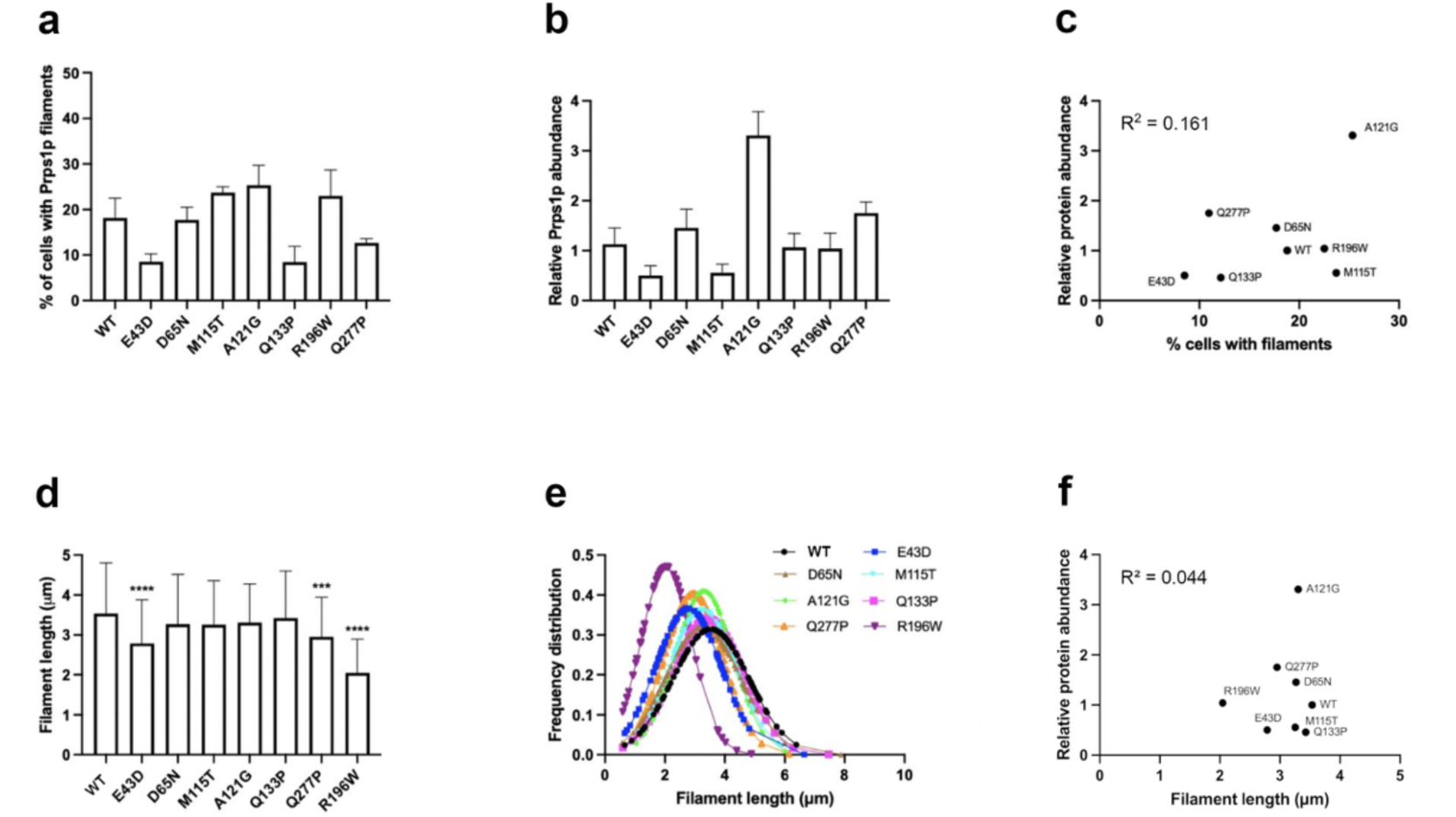
Loss-of-function PRPS1 variants differentially affect filament formation, protein abundance, and filament length in yeast. (a) Percentage of logarithmically growing yeast cells containing PRPS1 filaments. LOF variants displayed a broad range of filament formation phenotypes, with A121G and M115T forming filaments at higher frequencies than wild type, whereas E43D, Q133P, and Q277P formed filaments at reduced frequencies. (b) Relative PRPS1 abundance for the same strains, quantified as PRPS1/tubulin and normalized to wild-type levels. LOF variants exhibited diverse protein abundance profiles relative to wild type. (c) Correlation between filament formation frequency and PRPS1 protein abundance across LOF variants. No significant correlation was observed (R² = 0.161). (d) Mean filament lengths for wild-type and LOF PRPS1 variants. E43D, Q277P, and R196W showed significantly shorter filaments than wild type. (e) Kernel density distributions of filament lengths for wild-type and LOF variants. Wild-type filaments exhibited an approximately normal length distribution, whereas LOF variants displayed distinct filament-length distributions, most prominently for R196W. (f) Correlation between filament length and PRPS1 protein abundance across LOF variants. No significant correlation was observed (R² = 0.044). For filament-length measurements, 94 filaments per strain were analyzed across three independent biological replicates. Data are presented as mean ± SD where applicable. Filament-length distributions were compared using two-sided Mann–Whitney U tests comparing each variant with wild-type PRPS1. ***P < 0.001; ****P < 0.0001.

The D65N variant, associated with DFN2 (de Brouwer et al. 2010; Liu et al. 2010), accumulated at levels comparable to wild-type PRPS1 and formed filaments at similar frequencies (Fig. 3a and b). A121G, another mild hypomorphic allele associated with CMTX5 (Park et al. 2013), displayed a distinct phenotype: despite its classification as a mild LOF allele, A121G accumulated at approximately threefold higher levels than wild-type PRPS1 and formed filaments at significantly higher frequencies (Fig. 3a and b). Thus, the polymerization behavior of these variants does not simply parallel their classification based on catalytic impairment.

Yeast strains expressing the E43D or M115T variants, both associated with CMTX5, displayed approximately half the wild-type PRPS1 abundance. Whereas E43D formed filaments at proportionally lower frequencies, M115T formed filaments slightly more frequently than wild type (Fig. 3a and b), indicating that reduced protein abundance does not necessarily correspond to reduced filament formation. The CMTX5-associated R196W variant exhibited wild-type protein abundance and filament frequency but produced significantly shorter filaments with a markedly altered length distribution (Fig. 3d and e). R196W has previously been reported to exhibit extremely low enzymatic activity in patient cells (Al-Maawali et al. 2015).

Q133P and Q277P are strong LOF variants associated with severe PRPS1-deficiency phenotypes. Q133P is associated with Arts syndrome and markedly impaired catalytic activity (de Brouwer et al. 2007; de Brouwer et al. 2010), whereas Q277P has been associated with a CMTX5-Arts syndrome disease continuum and severe functional impairment (Synofzik et al. 2014). Q133P accumulated at near wild-type levels in yeast but formed approximately half as many filaments as wild-type PRPS1, with no detectable difference in filament length (Fig. 3a, d, and e). Q277P accumulated at approximately twice the wild-type level yet likewise formed approximately half as many filaments (Fig. 3a and b). These contrasting relationships between protein abundance and filament formation further indicate that steady-state PRPS1 levels alone do not determine polymerization behavior.

Across the LOF genotype panel, filament frequency showed no significant correlation with PRPS1 protein abundance (R² = 0.161; n = 8 genotypes; Fig. 3c). Analysis of filament length similarly revealed no significant correlation with protein abundance (R² = 0.044; n = 8 genotypes; Fig. 3f). E43D, Q277P, and R196W produced significantly shorter filaments than wild-type PRPS1 (Fig. 3d), and R196W displayed a particularly altered filament-length distribution (Fig. 3e). Together, these observations show that LOF variants can independently affect multiple measurable features of PRPS1 assembly, including filament frequency and length.

Overall, the diverse polymerization phenotypes observed among LOF variants are not explained by steady-state protein abundance alone. Together with the GOF analysis, these findings suggest that disease-associated substitutions influence PRPS1 assembly in an allele-specific manner, consistent with structural studies showing that PRPS1 filament architectures are associated with distinct allosteric and catalytic states (Hu et al. 2022; Hvorecny et al. 2023).

### PRPS1 filaments remain cytoplasmic in yeast regardless of variant class

PRPS1 filaments were restricted to the cytoplasm in yeast, consistent with previous studies describing cytoplasmic assemblies of metabolic enzymes (Noree et al. 2010; Noree et al. 2019). Because several disease-associated variants altered filament abundance and morphology, we next asked whether these phenotypes were accompanied by changes in subcellular localization. Analysis of Z-stack projections showed that PRPS1-positive structures remained cytoplasmic under all conditions examined and showed no detectable colocalization with the nuclear-envelope marker Nup157–GFP (Figure S3). Both GOF variants (D52H, D183H, A190V, V142L) and LOF variants (E43D, D65N, Q133P, Q277P) displayed similar localization patterns. Although A190V frequently formed sheet-like assemblies rather than elongated filaments, these structures also remained cytoplasmic. Thus, disease-associated variants alter PRPS1 filament abundance and morphology without detectably changing the overall subcellular localization of these assemblies in yeast.

### *PRPS1* variants are associated with altered actin organization in *prs3*Δ yeast

Because endogenous yeast PRS proteins have been linked to actin organization and cell-integrity pathways (Binley et al. 1999; Wang et al. 2004; Ugbogu et al. 2016), we next examined actin architecture in *prs3*Δ cells expressing human PRPS1. Wild-type yeast displayed the expected polarized cortical actin patches and organized actin cables, whereas *prs3*Δ cells frequently accumulated dense F-actin aggregates (Fig. 4a). Quantification confirmed that approximately half of *prs3*Δ cells contained F-actin aggregates, whereas such structures were rare in wild-type cells; expression of wild-type or disease-associated human PRPS1 in the *prs3*Δ background was also associated with a high frequency of actin aggregates (Fig. 4b).

**Figure 4.**
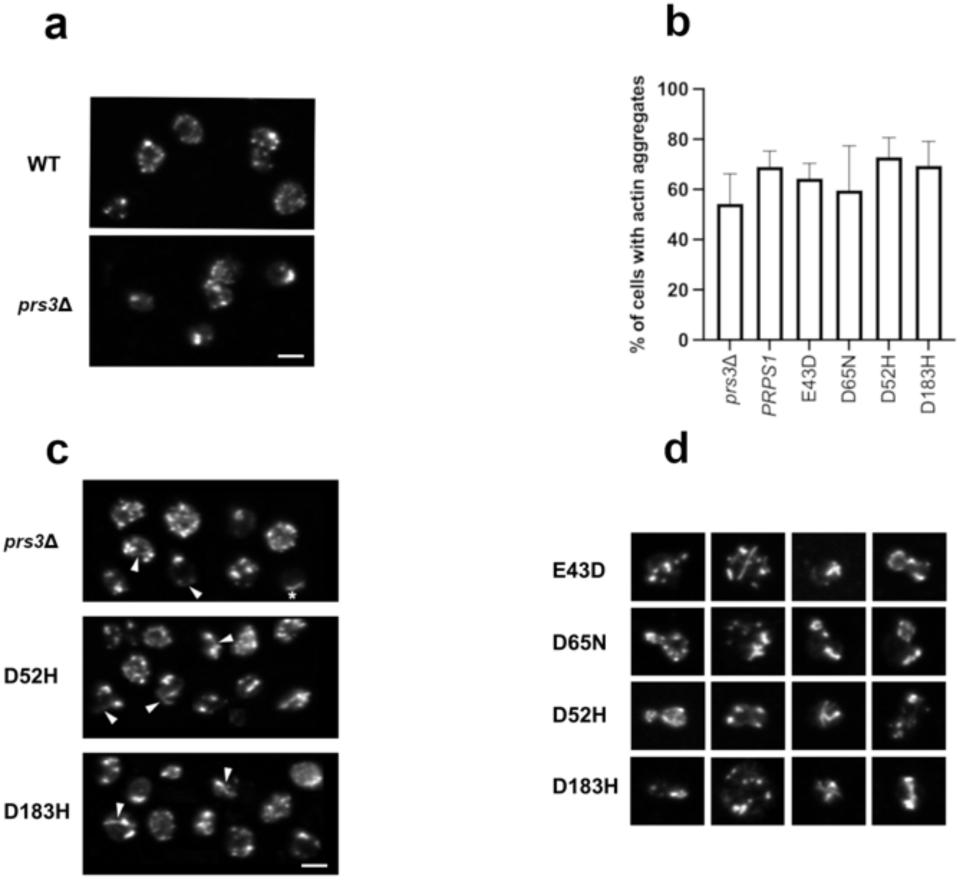
PRPS1 variants alter actin organization in *prs3*Δ yeast cells. (a) Representative images of F-actin organization in wild-type and *prs3*Δ cells after 24 h of growth. Wild-type cells display punctate cortical actin patches, whereas *prs3*Δ cells frequently exhibit dense F-actin aggregates. Scale bar, 5 µm. (b) Quantification of cells containing F-actin aggregates. Approximately half of *prs3*Δ cells exhibited F-actin aggregation, whereas PRPS1-expressing *prs3*Δ strains showed a high frequency of cells containing aggregates. More than 100 cells per strain were scored in each biological replicate. Data are presented as mean ± SD from 3–4 independent biological experiments. Comparisons between *prs3*Δ and each PRPS1-expressing strain were performed using two-sided unpaired t-tests; none of the comparisons reached P < 0.05. (c) Representative images of elongated cable-like actin structures observed in *prs3*Δ cells expressing PRPS1 variants. These structures were particularly apparent in cells expressing the gain-of-function variants D52H and D183H. Arrowheads indicate cable-like structures; an example not associated with a cortical patch is marked with an asterisk. Scale bar, 5 µm.(d) Additional actin abnormalities observed in post-diauxic *prs3*Δ cells expressing PRPS1 variants, including non-polarized cortical actin patches, misaligned or transverse actin bundles, and abnormal actin accumulation near the mother–bud neck.

In addition to these quantified aggregates, PRPS1-expressing *prs3*Δ cells displayed a range of more heterogeneous actin abnormalities, including elongated cable-like structures and altered cortical actin distributions. These phenotypes were observed with both GOF and LOF variants, although their morphology varied among alleles. GOF variants D52H and D183H frequently displayed prominent elongated actin structures extending across large portions of the cell cortex (Fig. 4c), whereas cells expressing the LOF variants E43D and D65N showed examples of disorganized cortical patches, transverse actin bundles, and abnormal actin accumulation near the mother–bud neck (Fig. 4d). These latter phenotypes were particularly apparent in post-diauxic cultures. Because of their morphological heterogeneity, these additional abnormalities were characterized qualitatively rather than used as a quantitative measure of actin disruption.

Together, these observations show that expression of both GOF and LOF PRPS1 variants in the *prs3*Δ background is associated with altered actin organization. The presence of related abnormalities across multiple disease-associated alleles supports an association between PRPS1 dysfunction and actin organization, although these experiments do not establish whether the relationship is direct or mediated through changes in cellular metabolism.

### Cable-like actin structures are more frequently observed with *PRPS1* gain-of-function alleles

In addition to F-actin aggregates, *prs3*Δ cells occasionally displayed elongated actin structures resembling cable-like assemblies previously described in actin-regulatory mutants (Belmont and Drubin 1998). These structures were observed in *prs3*Δ cells expressing wild-type or variant PRPS1 but appeared more frequently in cells expressing the GOF variants D52H and D183H (Fig. 4c). Because these structures were relatively infrequent and morphologically heterogeneous, they were assessed qualitatively, and their molecular composition remains unknown. Thus, their increased occurrence in D52H- and D183H-expressing cells identifies an additional actin phenotype associated with these variants but does not establish its underlying mechanism.

### *PRPS1* variants display diverse actin abnormalities in post-diauxic cells

Post-diauxic *prs3*Δ cells expressing GOF or LOF PRPS1 variants exhibited additional F-actin abnormalities, including non-polarized cortical actin patches in budding cells, misaligned or transverse actin bundles, and abnormal accumulation of actin near the mother–bud neck (Fig. 4d). Occasional cells also exhibited incomplete cytokinesis, although the basis of this phenotype remains unclear. These observations further illustrate the range of actin abnormalities associated with expression of disease-associated PRPS1 variants under post-diauxic conditions.

Representative examples are shown for LOF (E43D, D65N) and GOF (D52H, D183H) variants.

Images are representative of at least three independent biological replicates. The additional actin phenotypes shown in (c) and (d) were assessed qualitatively.

### *PRPS1* variants alter cell size distributions in prs3Δ yeast

Because actin organization influences cell size and growth control, we examined cell-size distributions in *prs3*Δ cells expressing wild-type or disease-associated PRPS1 variants. PRPS1-expressing *prs3*Δ strains showed increased variability in cell size relative to wild-type yeast (Fig. 5). Cells expressing the LOF variants E43D and D65N were enriched in smaller size classes, whereas cells expressing wild-type PRPS1 or the GOF variants D52H and D183H showed increased proportions of larger cells (Fig. 5a; Figure S4).

**Figure 5.**
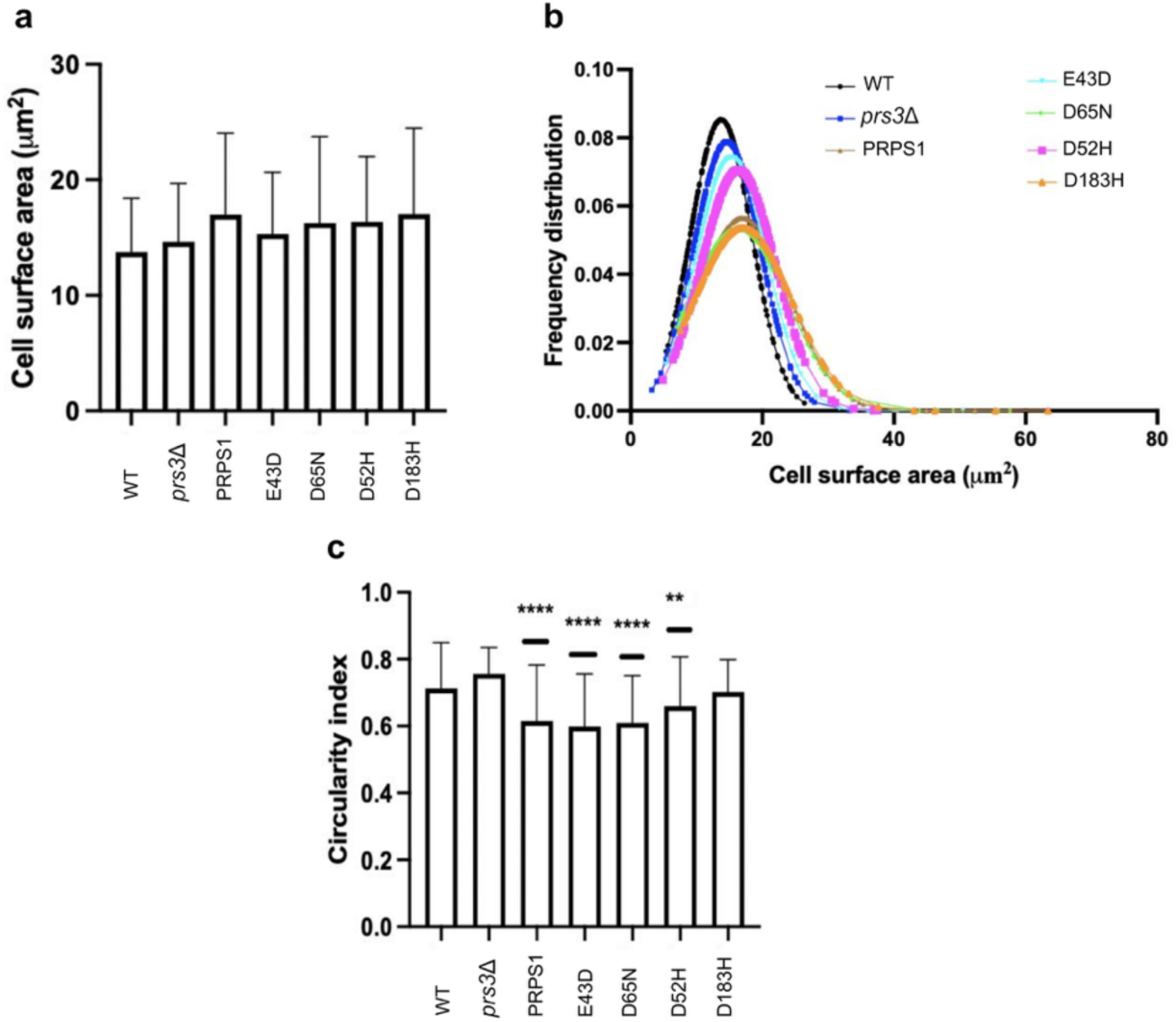
PRPS1 variants alter cell-size distributions and morphology in *prs3*Δ yeast. (a) Mean cell area (µm²) of unbudded wild-type, *prs3*Δ, and *prs3*Δ cells expressing wild-type PRPS1 or the indicated variants. Bars represent mean ± SD. (b) Kernel density estimation of cell-area distributions for the same strains, showing broader distributions in PRPS1-expressing *prs3*Δ backgrounds relative to wild type. For panels (a) and (b), 200 cells per strain were analyzed. (c) Circularity index (4π × area/perimeter²) for unbudded cells, where values approaching 1 indicate increased roundness. Cells expressing wild-type PRPS1, E43D, D65N, or D52H showed significantly reduced circularity relative to wild-type cells. Only unbudded cells were included in the analyses to avoid confounding effects of bud emergence. Statistical significance for circularity was assessed using a Kruskal–Wallis test followed by Dunn’s multiple-comparison test comparing each group with wild-type cells. For circularity analysis, 134–155 cells per strain were analyzed. **P < 0.01; ****P < 0.0001.

Kernel-density analysis revealed broader, flatter cell-size distributions across the PRPS1-expressing *prs3*Δ strains compared with wild type (Fig. 5b), indicating altered cell-size distributions under these conditions.

### *PRPS1* variants alter cell morphology

Because altered cell morphology has previously been associated with actin disorganization (Gross and Kinzy 2005; Gross and Kinzy 2007), we next asked whether the cytoskeletal abnormalities observed in PRPS1-expressing *prs3*Δ cells were accompanied by measurable changes in cell shape. We therefore quantified cell circularity, with higher values indicating more rounded cells.

Cells expressing wild-type PRPS1 or the E43D, D65N, and D52H variants exhibited significantly reduced circularity relative to wild-type cells (Kruskal–Wallis test with Dunn’s multiple-comparison correction, *P* < 0.0001; Fig. 5c). By contrast, *prs3*Δ cells alone did not show a detectable reduction in circularity relative to wild type. Thus, several PRPS1-expressing *prs3*Δ strains exhibited altered morphology relative to wild-type cells.

Together, the cell-size and circularity measurements show that expression of wild-type or disease-associated PRPS1 in the *prs3*Δ background is associated with changes in cellular morphology in addition to the actin phenotypes described above.

### Metabolic conditions associated with *ado1*Δ suppress PRPS1 filament formation

Because several superactivity PRPS1 variants formed abundant cytoplasmic filaments in yeast, we asked whether filament assembly was sensitive to a genetic background associated with altered nucleotide metabolism. Deletion of ADO1, which encodes adenosine kinase, has previously been shown to elevate intracellular adenosine and S-adenosylmethionine (SAM) levels (Kanai et al. 2013). We therefore expressed wild-type PRPS1 and the superactivity variants D52H and D183H in a prs3Δ ado1Δ background and quantified filament formation.

Compared with their behavior in *prs3*Δ cells, wild-type PRPS1, D52H, and D183H formed markedly fewer filaments in the *prs3*Δ *ado1*Δ background (Fig. 6a), showing that PRPS1 assembly is strongly altered under the cellular conditions associated with *ADO1* deletion.

**Figure 6.**
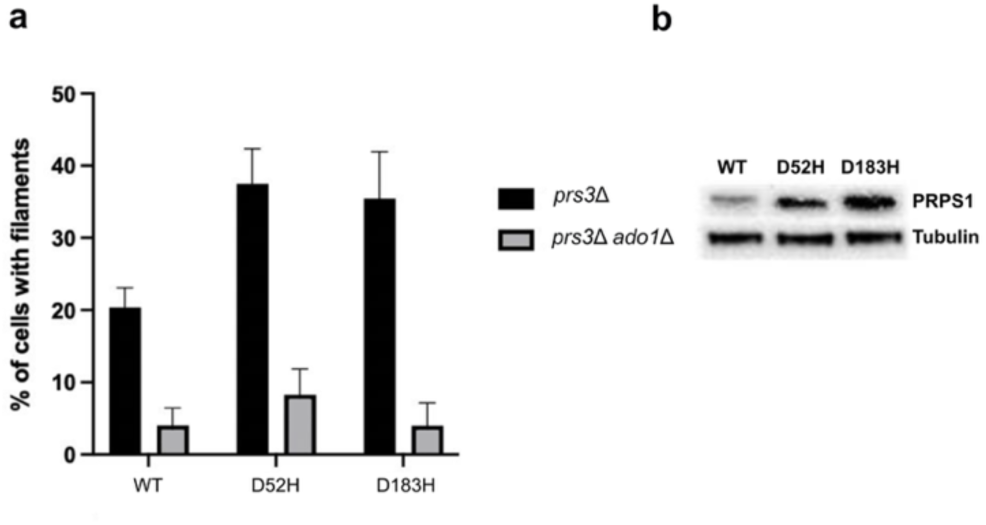
Metabolic conditions associated with *ado1*Δ suppress PRPS1 filament formation. (a) Percentage of *prs3*Δ and *prs3*Δ *ado1*Δ cells expressing wild-type, D52H, or D183H PRPS1 that contained cytoplasmic filaments. *ADO1* deletion reduced filament formation for all three PRPS1 proteins. Bars represent mean ± SD from at least three independent biological replicates. (b) Representative protein immunoblot showing PRPS1 abundance in *prs3*Δ *ado1*Δ cells expressing wild-type, D52H, or D183H PRPS1 variants. Tubulin served as a loading control. D52H and D183H remained highly expressed despite reduced filament formation in the *ado1*Δ background.

To determine whether reduced filament formation was accompanied by reduced PRPS1 abundance, we analyzed protein levels by immunoblotting. D52H and D183H remained highly abundant in *ado1*Δ cells, at approximately three- to fourfold higher levels than wild-type PRPS1 (Fig. 6b). Thus, the reduction in filament formation in the *ado1*Δ background cannot be attributed to decreased steady-state PRPS1 abundance.

Together, these findings indicate that PRPS1 filament formation is sensitive to the cellular context associated with *ADO1* deletion. Because we did not directly measure adenosine or SAM levels in these strains, these experiments do not establish which specific metabolic change is responsible for the reduction in polymerization. Previous work showing accumulation of adenosine and SAM in *ado1*Δ cells nevertheless identifies these metabolites as plausible contributors (Kanai et al. 2013).

### The D183H superactivity variant is associated with cytoplasmic PRPS1 polymerization in human fibroblasts

To determine whether disease-associated PRPS1 variants are accompanied by altered filament formation in human cells, we analyzed primary fibroblasts derived from two individuals carrying LOF variants (LOF1 and LOF2) and one individual carrying the feedback-resistant D183H superactivity variant (Maruyama Broyer 2016). LOF1 and LOF2 fibroblasts displayed markedly reduced PRPS1 protein levels (Fig. 7b), and PRPS1 filaments were rare in these cells (2.4% and 1.7%, respectively) compared with wild-type fibroblasts (22.2%; Fig. 7a).

**Figure 7.**
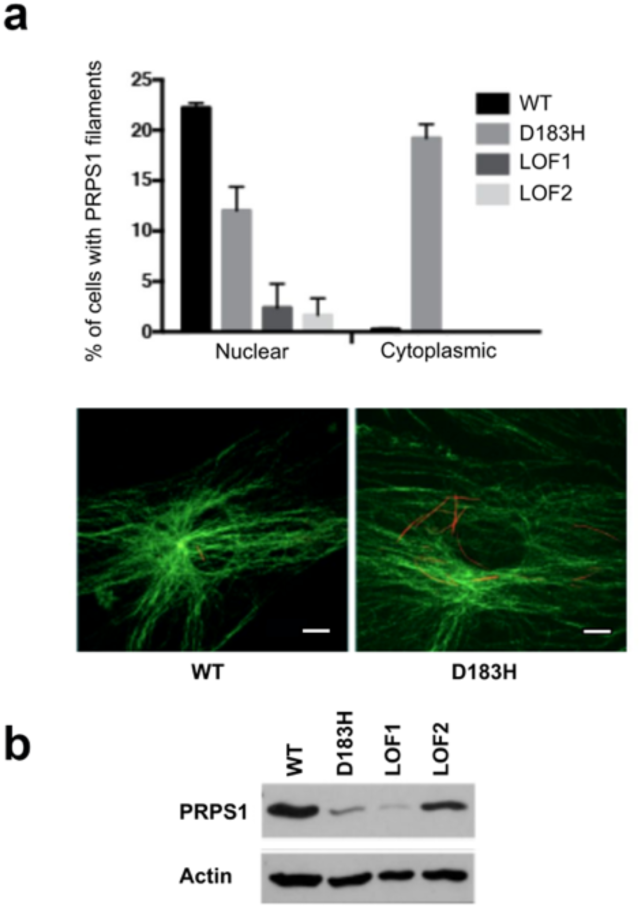
PRPS1 filament localization in human fibroblasts and cytoplasmic polymerization associated with the D183H superactivity variant. (a) Quantification of PRPS1 filament localization in primary human fibroblasts derived from wild-type (WT), D183H superactivity, and loss-of-function (LOF1, LOF2) individuals. WT fibroblasts exhibited nuclear PRPS1 filaments in 22.2% of cells and no detectable cytoplasmic filaments. D183H fibroblasts exhibited nuclear filaments in 12.0% of cells and cytoplasmic PRPS1 filaments in 19.2% of cells, whereas LOF fibroblasts showed markedly reduced nuclear filament formation (2.4% for LOF1; 1.7% for LOF2) and no detectable cytoplasmic filaments. Data represent counts aggregated from independent experiments. PRPS1 (red) and actin (green) were detected by immunofluorescence. Scale bars, 10 µm. (b) Protein immunoblot analysis of PRPS1 in WT, D183H, LOF1, and LOF2 fibroblasts. Actin served as a loading control.

D183H fibroblasts displayed a distinct localization pattern. Although overall PRPS1 abundance was not increased (Fig. 7b), 19.2% of D183H fibroblasts contained cytoplasmic PRPS1 filaments, whereas cytoplasmic filaments were not detected in wild-type or LOF fibroblasts (Fig. 7a). Nuclear filament frequency was lower in D183H fibroblasts (12.0%) than in wild-type cells (22.2%), indicating an altered subcellular distribution of PRPS1 assemblies in the D183H patient-derived line.

Cytoplasmic PRPS1 filaments in D183H fibroblasts did not visibly colocalize with actin structures. Together, these observations show that the D183H patient-derived fibroblast line is associated with an abnormal cytoplasmic PRPS1 polymerization phenotype not observed in the wild-type or LOF lines examined.

### *PRPS1* disease variants are associated with altered actin stress-fiber organization in patient-derived fibroblasts

*PRPS1*-related neurological disorders frequently include progressive sensorineural deafness and ataxia, phenotypes that have also been associated with defects in actin organization (Zhu et al. 2003; Rendtorff et al. 2006; Morín et al. 2009). We therefore examined actin stress-fiber organization in the patient-derived fibroblast lines.

At low confluency, PRPS1-variant fibroblasts retained partial actin polarity with only modest disorganization relative to wild-type cells, consistent with the corresponding quantitative analyses (Figure S5). As cells approached confluency, both LOF and D183H fibroblasts displayed pronounced loss of global actin alignment, with intersecting and circumferentially arranged stress fibers replacing the more ordered architecture observed in wild-type cells (Fig. 8). LOF2 fibroblasts showed particularly pronounced disorganization under both low- and high-confluency conditions.

**Figure 8.**
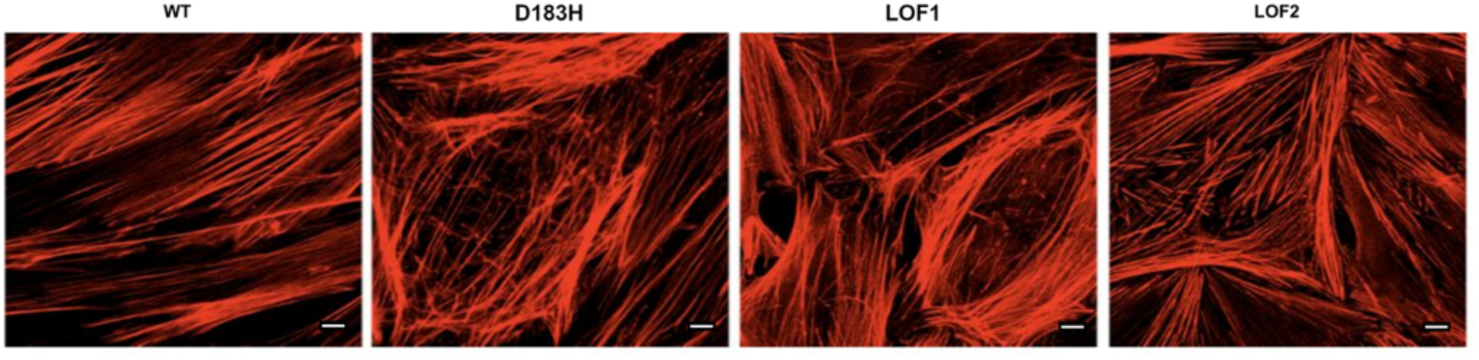
PRPS1 disease variants are associated with altered actin stress-fiber organization in patient-derived fibroblasts at high confluency. Wild-type (WT), D183H, and loss-of-function (LOF1 and LOF2) patient-derived fibroblasts were immunostained to visualize actin stress fibers at high confluency. At 9 days post-plating (100% confluency), WT fibroblasts exhibited highly polarized actin stress fibers aligned along a common cellular axis, whereas D183H, LOF1, and LOF2 fibroblasts displayed less organized stress-fiber arrangements, including intersecting and circumferentially oriented fibers. Representative images are shown. Scale bars, 10 µm.

Quantitative analysis showed broader stress-fiber angle distributions and reduced dominant orientation in the PRPS1-variant fibroblast lines, particularly at high confluency (Figure S5).

Within the set of patient-derived lines examined, both LOF and superactivity variants were therefore associated with altered actin organization. Together with the related phenotypes observed in yeast, these findings identify a similar association between PRPS1 dysfunction and altered cytoskeletal organization across the two experimental systems. Because the human-cell analysis comprises one superactivity and two LOF patient-derived lines, the extent to which these phenotypes generalize across additional PRPS1 variants remains to be established.

More than 45 cells per cell line were analyzed from randomized fields across three independent experiments. Quantitative analysis of actin stress-fiber orientation is presented in Figure S5.

## Discussion

PRPS1 encodes a central enzyme in nucleotide metabolism. Mutations in PRPS1 cause a wide spectrum of human disorders, including DFN2, CMTX5, Arts syndrome, and PRPS superactivity (Rosenberg and Chutorian 1967; Becker et al. 1988; Arts et al. 1993; de Brouwer et al. 2007; de Brouwer et al. 2010). Despite their opposing biochemical classifications as loss- and gain-of-function alleles, these mutations frequently produce overlapping neurological phenotypes, particularly sensorineural deafness and progressive ataxia. The molecular and cellular basis for this convergence remains unresolved. Our findings support a cellular framework in which PRPS1 polymerization is a metabolically responsive feature that varies among disease-associated alleles and is accompanied by changes in cellular organization.

An increasing number of metabolic enzymes are known to assemble into dynamic higher-order structures, ranging from filaments to metabolic condensates, that can influence pathway organization and activity (O’Connell et al. 2012; Ugbogu et al. 2016; Lynch et al. 2020). Such assemblies have been described across diverse systems, from yeast to human cells (Noree et al. 2010; Barry et al. 2014; Chang et al. 2018), while higher-order purine biosynthetic structures such as the human purinosome illustrate the broader importance of spatial organization in mammalian metabolism (An et al. 2008). Our findings place PRPS1 among this growing class of filament-forming metabolic enzymes and show that disease-associated substitutions alter its assembly behavior in allele-specific ways.

We first established that human PRPS1 is functionally active in *Saccharomyces cerevisiae*, rescuing the slow-growth phenotype of *prs1*Δ cells and supporting viability following *PRS5* deletion in a *prs3*Δ background. Nearly all disease-associated PRPS1 variants retained some ability to support viability, although rescue efficiency varied substantially among alleles. Q133P was the only variant unable to support viability, consistent with the severe functional impairment previously reported for this allele (de Brouwer et al. 2010). Importantly, complementation efficiency and polymerization phenotype did not map simply onto one another, indicating that these two cellular readouts capture different aspects of allele-specific PRPS1 behavior.

Recent structural studies provide a useful framework for interpreting the diversity of polymerization phenotypes observed here. Cryo-electron microscopy analyses have shown that human PRPS1 can form distinct filament architectures associated with different allosteric and catalytic states (Hvorecny et al. 2023). Our experiments did not directly measure PRPS1 conformation or enzymatic activity in different polymer states, but the allele-specific effects we observed are compatible with the idea that disease-associated substitutions influence higher-order assembly through distinct structural or regulatory mechanisms. D52H and D183H, for example, showed increased filament formation together with elevated protein abundance, whereas other GOF and LOF variants produced reduced filament frequency, altered filament length, or both. Across the variant panel, however, neither filament frequency nor filament length was simply predicted by steady-state protein abundance. Thus, PRPS1 assembly reflects more than protein accumulation alone, although the molecular determinants underlying these allele-specific differences remain to be established.

A second important finding is that PRPS1 polymerization is sensitive to cellular metabolic context. Deletion of *ADO1* strongly reduced filament formation by wild-type PRPS1 and by the hyperpolymerizing D52H and D183H variants despite continued high abundance of the variant proteins. Previous work has shown that *ado1*Δ cells accumulate adenosine and S-adenosylmethionine (SAM) (Kanai et al. 2013), but these metabolites were not measured directly in our experiments. We therefore cannot distinguish whether suppression of filament formation results specifically from elevated adenosine or SAM, from other metabolic consequences of *ADO1* deletion, or from a combination of these effects. Nevertheless, the *ado1*Δ phenotype demonstrates that PRPS1 assembly is sensitive to cellular metabolic context rather than being determined solely by protein abundance or variant class. Combined with prior evidence that nucleotide supplementation eliminates aberrant cytoplasmic PRPS1 filaments in D183H patient fibroblasts (Maruyama Broyer 2016), these findings further support the idea that PRPS1 assembly is sensitive to metabolic context. The ability of SAM and nicotinamide riboside supplementation to improve survival and metabolic function in PRPS1-deficient T cells further illustrates the broader sensitivity of PRPS1-associated phenotypes to metabolic intervention (Lenherr et al. 2021).

PRPS1 assembly also displayed distinct spatial organization in yeast and mammalian cells. In yeast, PRPS1 filaments remained cytoplasmic across the variants examined. In human fibroblasts, wild-type PRPS1 predominantly formed nuclear filaments, whereas the D183H patient-derived line displayed a prominent population of cytoplasmic filaments. Thus, the subcellular organization of PRPS1 assemblies differs between the two systems. Nevertheless, disease-associated variants altered PRPS1 assembly in both yeast and human cells, supporting the use of the two systems as complementary approaches for examining variant-dependent polymerization phenotypes.

A further observation emerging from both experimental systems is an association between PRPS1 dysfunction and altered actin organization. In yeast, *prs3*Δ cells displayed increased F-actin aggregation, and PRPS1-expressing strains showed additional allele-dependent abnormalities in actin organization, cell-size distributions, and morphology. In patient-derived fibroblasts, both LOF and D183H lines showed altered stress-fiber organization, particularly at high confluency. Actin dynamics are themselves sensitive to nucleotide state and cellular metabolism (DeWane et al. 2021; Ali et al. 2022; Liu et al. 2022), providing several plausible routes through which PRPS1 dysfunction could influence cytoskeletal organization. However, our experiments do not distinguish between a direct connection involving PRPS1 assembly, an indirect consequence of altered metabolism or nucleotide pools, or parallel consequences of PRPS1 dysfunction. The previously reported disruption of the lens-fiber actin network following *prps1a* truncation in zebrafish (Begovich et al. 2020) nevertheless provides independent support for a broader relationship between PRPS1 function and cytoskeletal organization.

One particularly distinctive phenotype was observed in fibroblasts carrying the D183H superactivity variant. Whereas cytoplasmic PRPS1 filaments were not detected in the wild-type or LOF lines examined, they were present in approximately 19% of D183H cells. These structures resemble the increased assembly phenotype observed for D183H in yeast, although their subcellular localization differs between the two systems. Their functional significance remains unknown, and the absence of visible colocalization with actin does not establish whether they directly contribute to the cytoskeletal phenotype. Nevertheless, the D183H fibroblast phenotype identifies altered spatial organization of PRPS1 assembly as a prominent cellular feature associated with this superactivity variant.

Several limitations define the interpretation of these findings. We did not manipulate PRPS1 polymerization independently of catalytic state and therefore cannot establish whether altered polymerization directly causes the actin or morphological phenotypes observed here. Similarly, enzymatic activity, PRPP, or nucleotide pools were not measured across different polymerization states, and the *ado1*Δ experiments use a genetic metabolic perturbation rather than direct manipulation of individual metabolites. Some of the heterogeneous yeast actin phenotypes were assessed qualitatively, although F-actin aggregation was quantified, and the human-cell analysis was restricted to one D183H superactivity line and two independent LOF patient-derived lines. Additional patient-derived or isogenic models will therefore be required to determine how broadly these phenotypes extend across PRPS1 disease alleles. These limitations do not alter the observed relationships among genotype, polymerization, metabolic context, and cellular organization, but they constrain conclusions regarding the underlying mechanism.

Together, our findings establish that disease-associated PRPS1 variants produce distinct higher-order assembly phenotypes that are not simply predicted by steady-state protein abundance, that PRPS1 filament formation is sensitive to metabolic context, and that functionally distinct PRPS1 alleles are associated with altered actin organization in both yeast and human cells. Rather than establishing polymerization as a causal regulatory mechanism, these results identify it as a metabolically responsive cellular feature associated with PRPS1 functional state and cytoskeletal organization. This framework provides a basis for future experiments aimed at determining how PRPS1 assembly, enzymatic activity, metabolic flux, and cytoskeletal organization are mechanistically connected and how these relationships may contribute to the overlapping phenotypes of PRPS1-associated disorders.

## Acknowledgments

We thank Alec Lee for permission to use filament quantification data, Dr. Lorraine Pillus and Dr. Ana Lilia Torres-Machorro for critical reading of the manuscript, and the late Dr. William L. Nyhan for providing the patient-derived fibroblast line carrying the D183H PRPS1 variant, originally described in Nyhan et al. (1969).

ChatGPT (OpenAI; GPT-5.6 Sol, ChatGPT interface) was used through iterative natural-language prompts requesting language editing, text organization, and consistency checks. All AI-assisted output was reviewed and, where appropriate, edited by the authors, who take responsibility for the final content of the manuscript.

