## Supplemental Data 1 for "Functional analysis of *PRPS1* disease alleles reveals altered polymerization and cytoskeletal organization"

**This PDF file includes:**

Tables S1 and S2

Supplementary Figures S1–S5

**Table S1. Structural and functional classification of *PRPS1* alleles analyzed in this study.**

All PRPS1 missense variants analyzed in this study have been previously described in the literature. Structural locations, functional classifications, and proposed mechanistic effects were compiled from prior biochemical, structural, and clinical studies to facilitate interpretation of the experimental data. Functional classes reflect previously published classifications and were not assigned in this work. References supporting each allele classification are provided in the table.

| Structural region | Variant(s) | Functional class | Mechanistic effects | References |
|---|---|---|---|---|
| Allosteric site I | **D52H** | Gain-of-function | Perturbs allosteric site I and reduces ADP-mediated feedback inhibition | de Brouwer et al. 2010; Chen et al. 2015 |
| Allosteric site II | **L129I** | Gain-of-function | Perturbs local structure at allosteric site II, impairing feedback regulation | de Brouwer et al. 2010 |
| Homodimer interface | **H193Q** | Gain-of-function | Disrupts intersubunit interactions and perturbs allosteric regulation | de Brouwer et al. 2010; Chen et al. 2015 |
| Homodimer/hexamer interface | **N114S** | Gain-of-function | Disrupts interface-stabilizing interactions and weakens ATP-induced higher-order assembly | Liu et al. 2009; de Brouwer et al. 2010 |
| Homodimer interface / β9–β10 region | **D183H** | Gain-of-function | Disrupts intersubunit interactions and perturbs allosteric regulation | de Brouwer et al. 2010; Chen et al. 2015 |
| Homodimer interface | **A190V** | Gain-of-function | Disturbs hydrophobic packing at the dimer interface and perturbs allosteric regulation | de Brouwer et al. 2010; Chen et al. 2015 |
| Trimer interface / ATP-binding region | **E43D** | Loss-of-function | Predicted to perturb ATP binding and allosteric regulation | Kim et al. 2007; de Brouwer et al. 2010 |
| Allosteric/ATP-binding region | **V142L** | Mixed phenotype | Predicted to affect allosteric inhibition and ATP binding, producing combined gain- and loss-of-function features | Moran et al. 2012 |
| ATP-binding region | **D65N** | Loss-of-function | Perturbs the ATP-binding β2–β3 region and reduces PRPS1 activity | Liu et al. 2010; Chen et al. 2015 |

| Structural region | Variant(s) | Functional class | Mechanistic effects | References |
|---|---|---|---|---|
| C-terminal catalytic region near R5P-binding site | **Q277P** | Loss-of-function | Destabilizes local structure and the region surrounding the R5P-binding site, severely impairing activity | Synofzik et al. 2014 |
| Inter-subunit interaction / catalytic-site region | **A121G** | Loss-of-function | Predicted to affect higher-order subunit interactions and/or the catalytic site | Park et al. 2013 |
| Homodimer interface | **M115T** | Loss-of-function | Reduces hydrophobic interactions at the dimer interface and decreases PRPS1 activity | Kim et al. 2007; Chen et al. 2015 |
| Homodimer interface / catalytic-loop–ATP-binding region | **Q133P** | Severe loss-of-function | Disrupts dimer-interface interactions and destabilizes catalytic-loop/ATP-binding-site organization | de Brouwer et al. 2007; Chen et al. 2015 |
| Catalytic loop / R5P-contacting region | **R196W** | Loss-of-function | Severely reduces PRPS1 activity; R196 is a catalytic-loop residue that contacts R5P | Al-Maawali et al. 2015; Hvorecny et al. 2023 |

**Table S2. Growth rescue spectrum of PRPS1 disease variants in the *prs3*Δ *prs5*Δ yeast background.**

Growth rescue phenotypes were assigned based on serial dilution assays of *prs3*Δ *prs5*Δ yeast strains expressing plasmid-borne human PRPS1 variants (Figure 1b). "Strong," "Intermediate," "Weak," and "None" reflect the relative ability of each variant to restore growth compared with wild-type PRPS1 under identical conditions and are intended as qualitative descriptors rather than quantitative measurements.

| *PRPS1* variant | Functional classification | Yeast growth rescue |
|---|---|---|
| **D52H** | Gain-of-function | Strong |
| **A190V** | Gain-of-function | Strong |
| **D183H** | Gain-of-function | Intermediate |
| **N114S** | Gain-of-function | Intermediate |
| **V142L** | Mixed phenotype | Weak |
| **L129I** | Gain-of-function | Weak |
| **H193Q** | Gain-of-function | Weak |
| **R196W** | Loss-of-function | Strong |
| **M115T** | Loss-of-function | Intermediate |
| **A121G** | Loss-of-function | Weak |
| **Q277P** | Loss-of-function | Weak |
| **D65N** | Loss-of-function | Weak |
| **E43D** | Loss-of-function | Weak |
| **Q133P** | Loss-of-function | None |

**Figures**

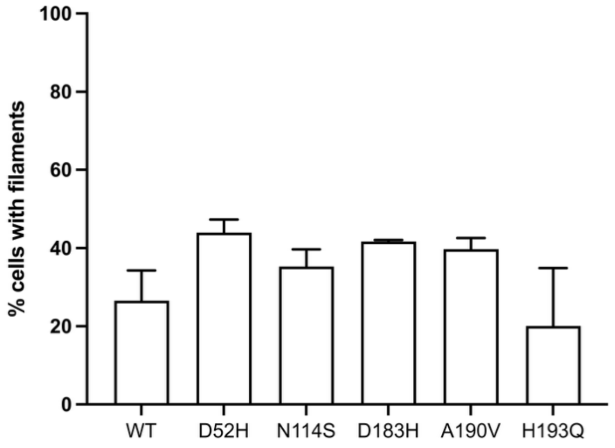


**Figure S1. PRPS1 filament formation in a wild-type yeast background.** The percentage of cells containing PRPS1 filaments was quantified for wild-type PRPS1 and gain-of-function variants (D52H, N114S, D183H, A190V, and H193Q) expressed in a wild-type genetic background. These experiments were performed independently of the *prs3*Δ experimental series presented in the main figures and are therefore shown separately. Bars represent mean ± SD from 3–4 independent biological replicates. Protein abundance was not assessed in this experiment.

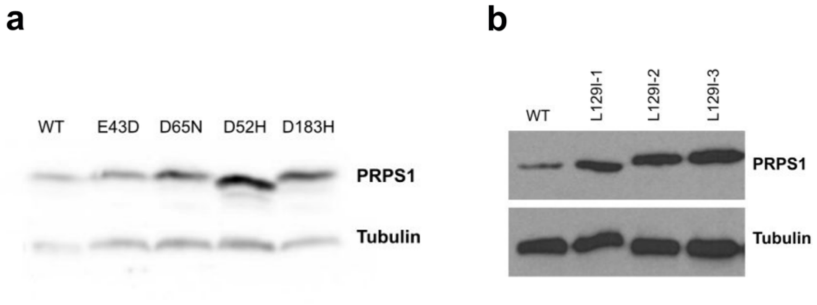


**Figure S2. Representative immunoblots showing PRPS1 protein abundance.** (a) Immunoblot showing wild-type PRPS1 and selected gain- and loss-of-function variants (E43D, D65N, D52H, and D183H) quantified in Figs. 2b and 3b.

(b) Independent immunoblot showing PRPS1 L129I abundance relative to wild type.

Whole-cell extracts from logarithmically growing *prs3*Δ cells expressing wild-type or variant PRPS1 were probed with anti-PRPS1 antibody; tubulin served as a loading control. Blots are representative of the experiments used for quantitative analyses shown in the main figures, with band intensities quantified across independent biological replicates.

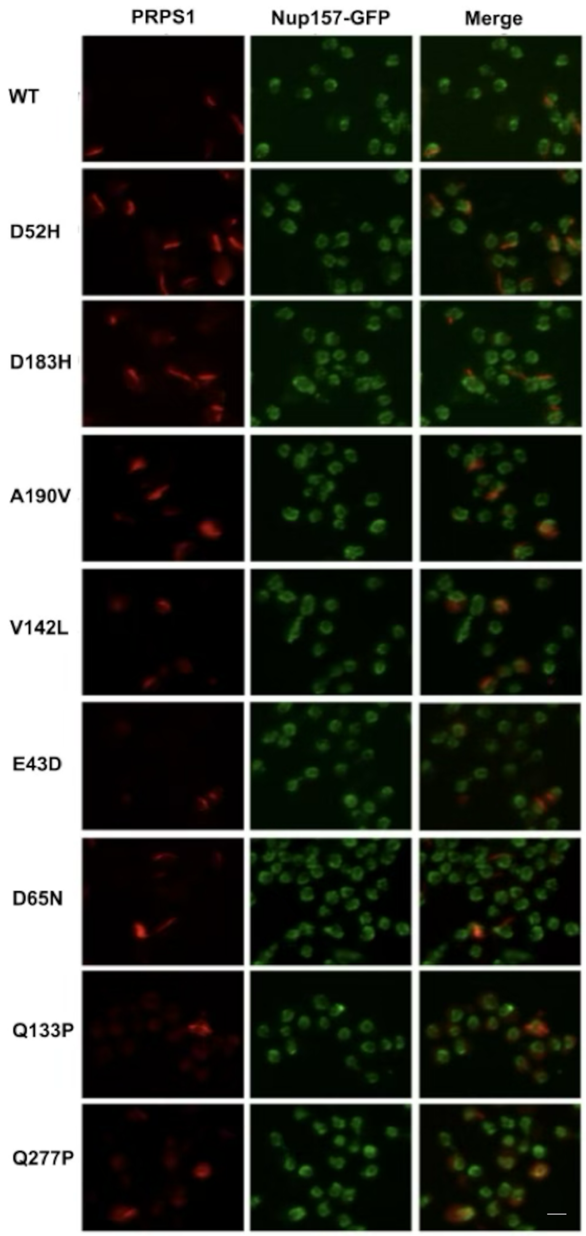


**Figure S3. PRPS1 filaments do not localize to the nuclear envelope in yeast.** Yeast cells expressing wild-type or variant PRPS1 were immunostained for PRPS1 (red), and the nuclear envelope marker Nup157–GFP was visualized in green. In all strains examined, PRPS1 structures remained cytoplasmic and showed no detectable colocalization with the nuclear envelope. For A190V, a representative image containing sheet-like PRPS1 assemblies is shown; these structures were excluded from filament quantification. Scale bar, 5 μm.

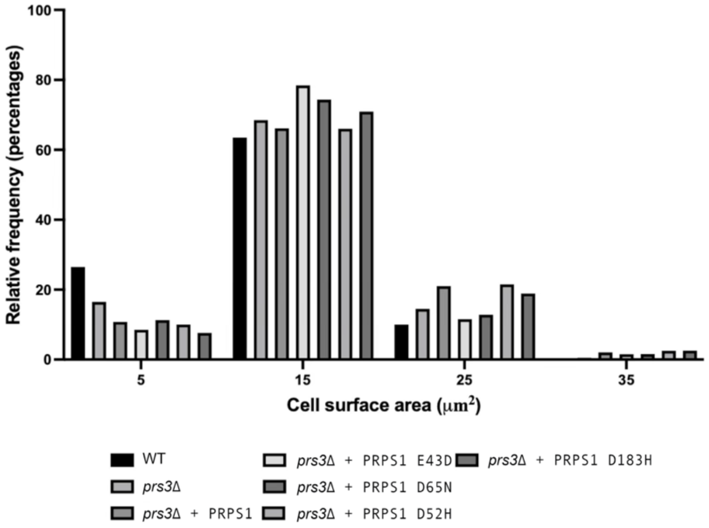


**Figure S4. Cell area distributions of *prs3*Δ strains expressing PRPS1 variants.** Relative frequency histograms of cell area for unbudded wild-type cells, *prs3*Δ cells, and *prs3*Δ cells expressing wild-type PRPS1 or the indicated variants. PRPS1-expressing *prs3*Δ strains displayed broader cell-area distributions relative to wild type. Data correspond to the same cell populations quantified in Figure 5a and analyzed by kernel density estimation in Figure 5b.

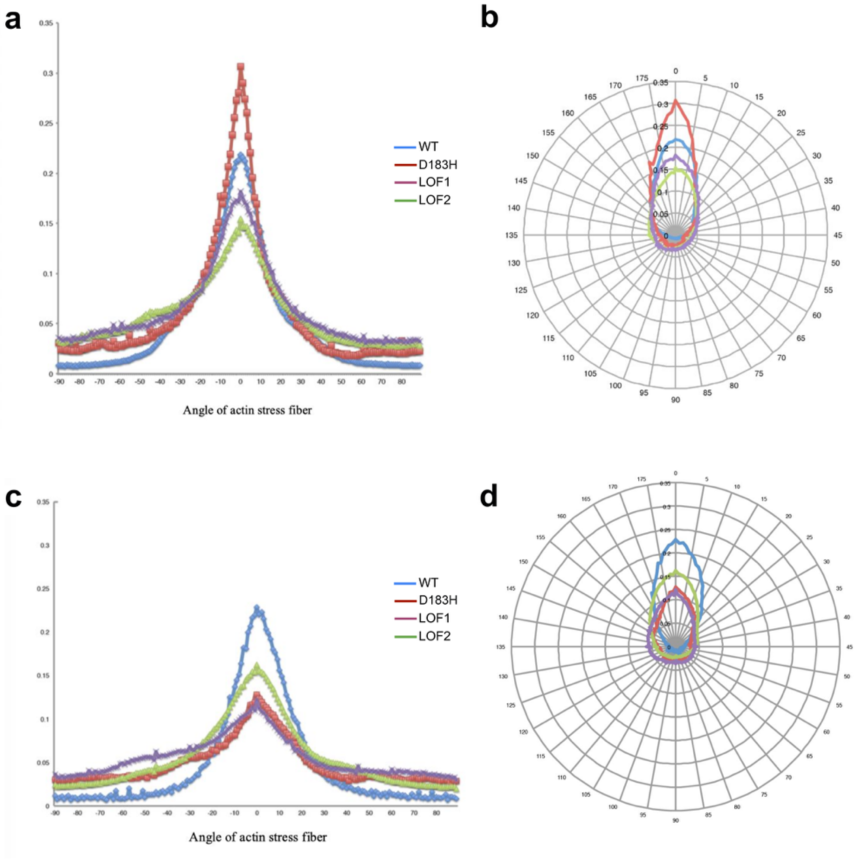


**Figure S5. Quantification of actin stress-fiber orientation in human fibroblasts.** (a) Distribution of actin stress-fiber angles relative to the dominant cellular polarity axis in wild-type (WT), D183H superactivity, and loss-of-function (LOF1 and LOF2) fibroblasts at 50% confluency.

(b) Polar plot of stress-fiber orientation at 50% confluency.

(c) Distribution of actin stress-fiber angles at 100% confluency.

(d) Polar plot of stress-fiber orientation at 100% confluency.

Actin fibers were segmented and their orientation quantified using identical image-processing parameters across all conditions. WT fibroblasts exhibited relatively narrow angular distributions consistent with coordinated stress-fiber alignment, whereas D183H and LOF fibroblasts displayed broader, less polarized distributions, particularly at high confluency. More than 45 cells per cell line were analyzed from randomized fields across three independent biological experiments.
